# Pseudohypoxic HIF pathway activation dysregulates collagen structure-function in human lung fibrosis

**DOI:** 10.1101/2021.05.11.443615

**Authors:** Christopher Brereton, Liudi Yao, Yilu Zhou, Milica Vukmirovic, Joseph Bell, Robert A. Ridley, Elizabeth R. Davies, Lareb S.N. Dean, Orestis G. Andriotis, Franco Conforti, Soran Mohammed, Tim Wallis, Ali Tavassoli, R. Ewing, Aiman Alzetani, Ben G. Marshall, Sophie V. Fletcher, Phillipp J. Thurner, Aurelie Fabre, Naftali Kaminski, Luca Richeldi, Atul Bhaskar, Matthew Loxham, Donna E. Davies, Yihua Wang, Mark G. Jones

**Author notes:** These authors contributed equally to this work (Christopher Brereton, Liudi Yao). Correspondence should be addressed to MGJ or YW.

## Abstract

Extracellular matrix (ECM) stiffening with downstream activation of mechanosensitive pathways is strongly implicated in fibrosis. We previously reported that altered collagen nanoarchitecture is a key determinant of pathogenetic ECM structure-function in human fibrosis (Jones et al., 2018). Here, through human tissue, bioinformatic and *ex vivo* studies we show that hypoxia-inducible factor (HIF) pathway activation is a critical pathway for this process regardless of oxygen status (pseudohypoxia). Whilst TGFβ increased rate of fibrillar collagen synthesis, HIF pathway activation was required to dysregulate post-translational modification of fibrillar collagen, promoting ‘bone-type’ cross-linking, altering collagen nanostructure, and increasing tissue stiffness. *In vitro*, knock down of Factor Inhibiting HIF (FIH) or oxidative stress caused pseudohypoxic HIF activation in normal fibroblasts. In contrast, endogenous FIH activity was reduced in fibroblasts from patients with lung fibrosis in association with significantly increased normoxic HIF pathway activation. In human lung fibrosis tissue, HIF mediated signalling was increased at sites of active fibrogenesis whilst subpopulations of IPF lung mesenchymal cells had increases in both HIF and oxidative stress scores. Our data demonstrate that oxidative stress can drive pseudohypoxic HIF pathway activation which is a critical regulator of pathogenetic collagen structure-function in fibrosis.

## Introduction

We previously identified that in the lung tissue of patients with idiopathic pulmonary fibrosis (IPF) there is increased ‘bone-type’ pyridinoline collagen cross-linking and altered collagen fibril nano-architecture, with individual collagen fibrils being structurally and functionally abnormal^1^. This was associated with increased tissue expression of lysyl hydroxylase 2 (LH2/PLOD2, which catalyses telopeptide lysine hydroxylation to determine pyridinoline cross-linking) and the lysyl oxidase-like (LOXL) enzymes LOXL2 and LOXL3, which initiate covalent collagen cross-linking^1^. This ‘bone-type’ pyridinoline cross-linking, rather than any change in collagen deposition per se, determined increased IPF tissue stiffness. Inhibiting ‘bone-type’ cross-linking normalised mechano-homeostasis and limited the self-sustaining effects of ECM on fibrosis progression. Whilst identifying the importance of altered collagen nanoarchitecture to human lung fibrosis pathogenesis, the upstream mechanisms that dysregulate collagen structure-function to promote progressive fibrosis rather than tissue repair were not determined. Here, we investigated possible mechanisms and established their relevance to human lung fibrosis.

## Results

### The ‘bone-type’ collagen fibrillogenesis genes PLOD2 and LOXL2 are co-expressed at sites of active fibrogenesis

In our previous work comparing human IPF lung tissue with age-matched control lung tissue, we identified that in bulk IPF lung tissue lysates there are significant increases in the relative expression of the collagen modifying enzymes *LOXL2, LOXL3*, and *LOXL4*, as well as *PLOD2* (also known as lysyl hydroxylase or LH2)^1^. To further investigate this observation, we firstly studied the transcriptomic profiles of fibroblast foci, the sites of active fibrogenesis in IPF. We analysed a data set we recently generated by integrating laser-capture-microdissection and RNA-Seq (LCMD/RNA-seq) which enabled profiling of the *in situ* transcriptome of fibroblast foci as well as alveolar septae from control tissue and IPF tissue (Gene Expression Omnibus (GEO) GSE169500). The LOXL enzyme with the greatest expression in fibroblast foci was *LOXL2* (Fig. 1a-e). Whilst *LOXL3* and *LOXL4* expression was increased within fibroblast foci, only limited expression was identified (Fig. 1d and e). Furthermore, *PLOD2* expression was significantly increased within fibroblast foci (Fig. 1f), and *PLOD2* expression correlated (r=0.63, p=0.04) with *LOXL2* (Fig. 1g) but not other LOXL enzymes (Supplementary Figure 1a-d), suggesting possible co-ordinated regulation of *PLOD2* and *LOXL2* gene expression. In contrast, expression of the major collagen fibrillogenesis gene *COL1A1* did not significantly correlate with their expression (Supplementary Fig. 1e), suggesting that, in lung fibrosis, distinct pathways might promote pyridinoline cross-linking to dysregulate collagen fibril nano-structure independently of pathways regulating major fibrillar collagen synthesis. We then performed RNA *in situ* hybridisation upon IPF lung tissue, confirming that the greatest expression of *LOXL2* and *PLOD2* in IPF tissue was by mesenchymal cells within fibroblast foci, and that *LOXL2* and *PLOD2* were co-expressed within the same cells (Fig. 1h).

**Figure 1.**
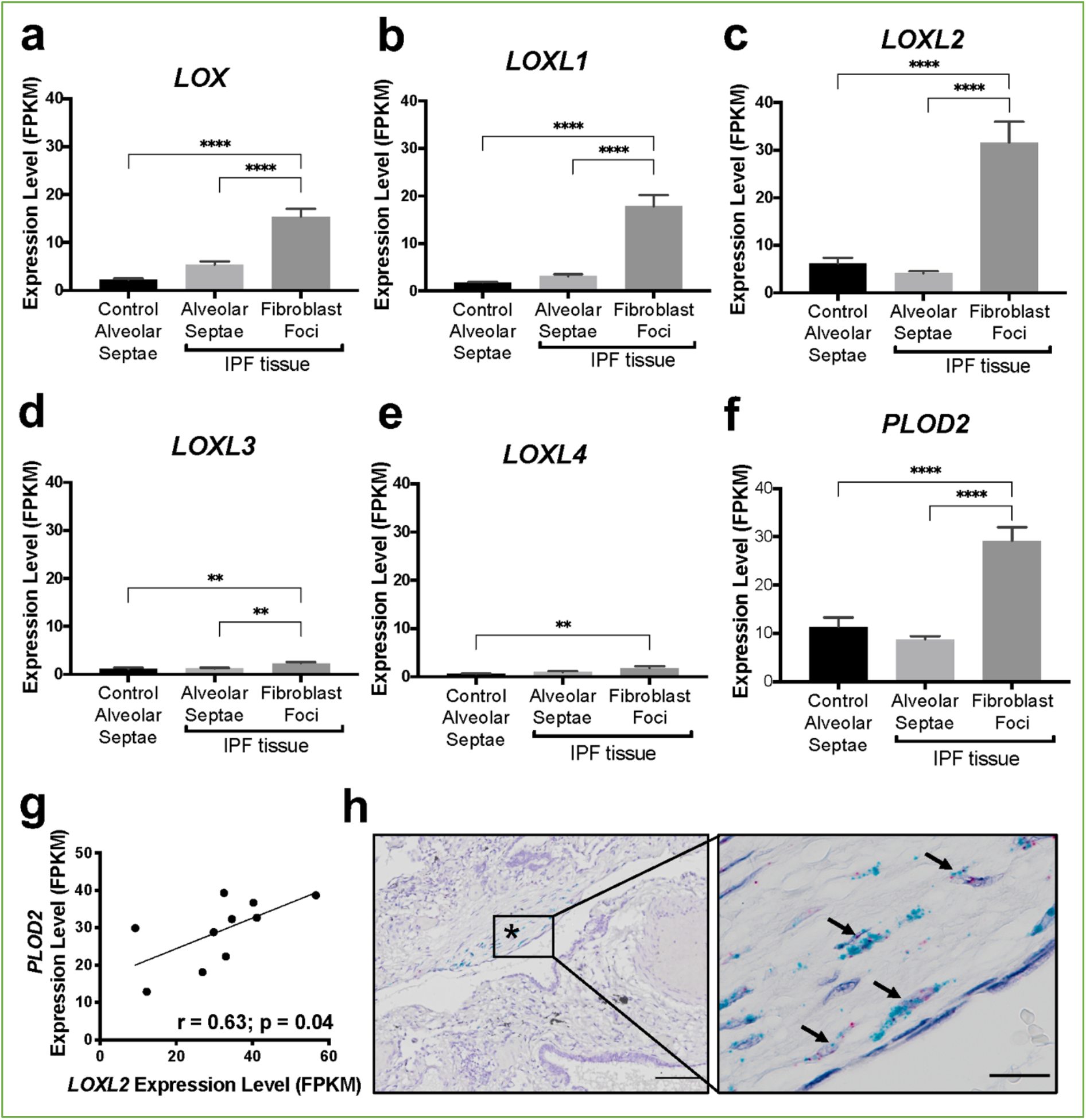
The collagen cross-linking enzymes PLOD2 and LOXL2 are co-expressed at sites of active fibrogenesis in IPF. (A-F) Expression of *LOX, LOXL1, LOXL2, LOXL3, LOXL4* and *PLOD2* in healthy alveolar septae, IPF alveolar septae and IPF fibroblast foci (n= 10 individual healthy and IPF donors). Relative expression levels are calculated as Fragments Per Kilobase of transcript per Million mapped reads (FPKM). Bars represent standard geometric means. *p<0.05; **p<0.01; ***p<0.001; ****p<0.0001 by Tukey’s multiple comparisons test. (G) Scatterplot of paired fibroblast foci data from (C) and (F) were plotted to compare expression of PLOD2 and LOXL2 (Spearman correlation coefficient r=0.63, p=0.04). (H) Representative mRNA expression of *PLOD2* (red chromagen) and *LOXL2* (green chromagen) in IPF lung tissue using RNAscope^®^ RNA in-situ hybridisation. A fibroblastic focus is identified by * and arrows identify co-expression pattern. Left scale bar 100 μm, right scale bar 20 μm.

### HIF pathway activation is a key inducer of PLOD2 and LOXL2 expression in lung fibroblasts

To investigate common regulators of *PLOD2* and *LOXL2* in lung fibrosis, we studied their expression in primary human lung fibroblasts over a 72 hour time course following activation (Supplementary Fig. 2a) of transforming growth factor beta (TGFβ), epidermal growth factor (EGF), hypoxia inducible factors (HIF) or Wnt signalling pathways, each of which have been implicated in fibrogenesis^2-7^. The hypoxia mimetic dimethyloxallylglycine (DMOG)^8^, a prolyl hydroxylase inhibitor that stabilises HIF1*α* and HIF2*α*, most strongly upregulated both PLOD2 and LOXL2 gene and protein expression (Fig. 2a-c) but did not induce expression of interstitial collagen genes (*COL1A1, COL3A1*) (Fig. 2d and Supplementary Fig. 2b). In contrast, TGFβ1 strongly induced *COL1A1* and *COL3A1* and this was associated with a smaller up-regulation of *PLOD2* at 24 hours and of *LOXL2* at 72 hours (Fig. 2a-d; Supplementary Fig. 2b). No induction of *PLOD2* or *LOXL2* was identified with canonical Wnt (Wnt3a), non-canonical Wnt (Wnt5a) or EGF pathway activation (Fig. 2a-c). We further extended these observations by showing that treatment with the HIF*α* selective prolyl 2-hydroxylase inhibitor, N-[[1,2-Dihydro-4-hydroxy-2-oxo-1-(phenylmethyl)-3-quinolinyl]carbonyl]-glycine (IOX2)^8^ or culture for 24h under hypoxic conditions induced expression of *PLOD2* and *LOXL2* (Supplementary Fig. 2c-d), with immunofluorescent staining confirming an increase in intracellular LOXL2 expression following DMOG or IOX2 treatment in comparison to treatment with TGFβ1 (Fig. 2e).

**Figure 2.**
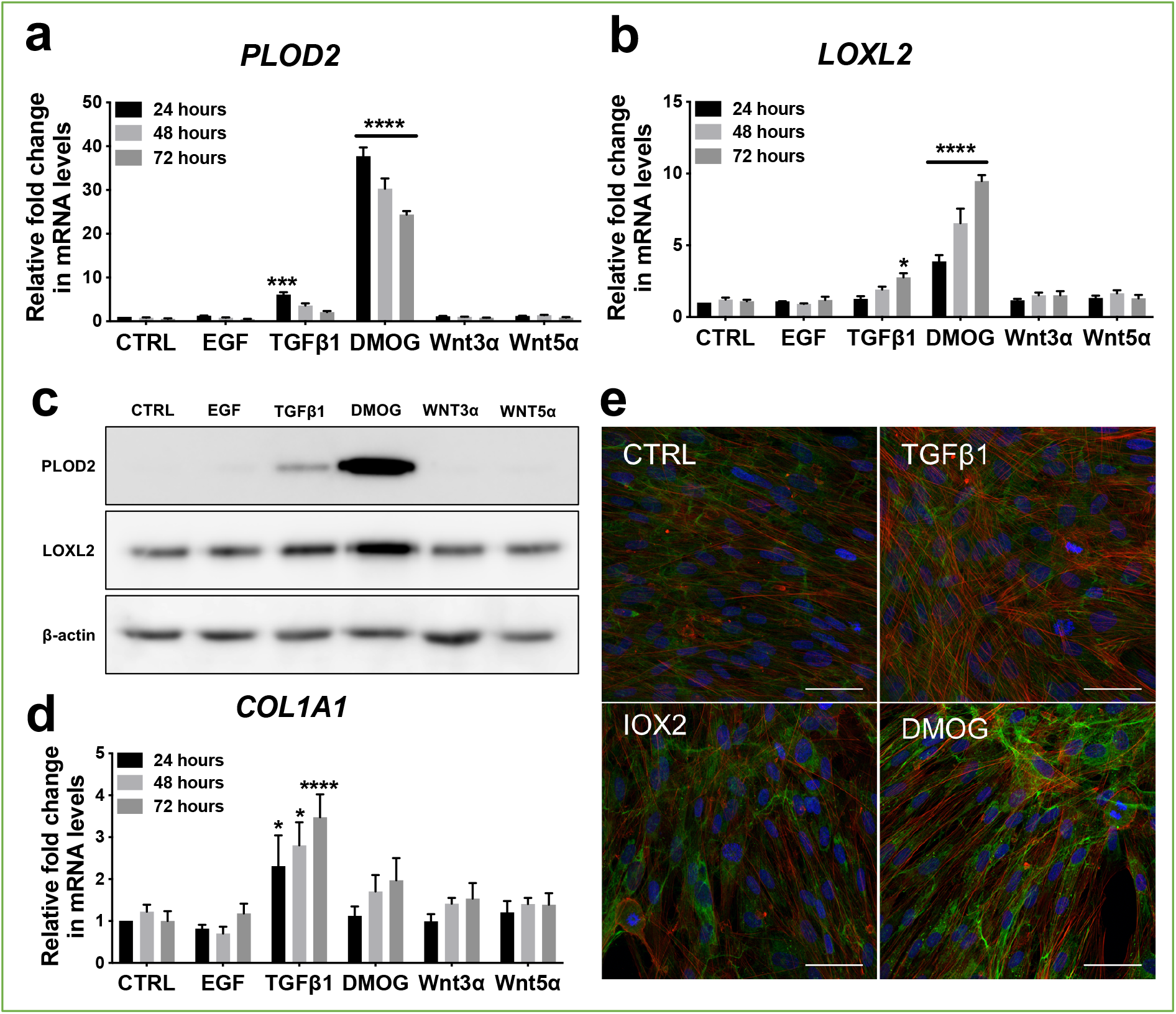
Hypoxia mimetics strongly promote PLOD2 and LOXL2 expression in lung fibroblasts. (A-B, D) Relative gene expression using the ΔΔCt method of *PLOD2, LOXL2* and *COL1A1* in healthy lung fibroblasts over a 72 hour time course in the presence of EGF, TGFβ1, the hypoxia mimetic DMOG, Wnt3a, Wnt5a, or vehicle control. n=3 independent experiments. Bars indicate geometric means. *p<0.05; ***p<0.001; ****p<0.0001 by Dunnett’s multiple comparisons test. (C) Protein expression of PLOD2 and LOXL2 at 72 hours. β-actin loading control. The full blots are shown in Figure 2-source data 1. (E) Representative immunofluorescence images of healthy lung fibroblasts fixed after 24 hr and stained for LOXL2 (green), phalloidin (red) and DAPI (blue). Scale bar 50 μm.

Transcriptional activation of HIF pathways requires assembly of a heterodimer between HIF1*α* or HIF2*α* and their obligate binding partner HIF1*β*^9, 10^. To confirm the dependence of the induction of PLOD2 and LOXL2 expression upon HIF pathways, siRNA knockdown against HIF1*α*, HIF2*α*, and HIF1β was performed (Fig. 3a). The knockdown of HIF1*α*, but not HIF2*α* prevented DMOG induction of PLOD2 mRNA and protein expression, whilst LOXL2 required silencing of both HIF1*α* and HIF2*α* or HIF1β (Fig. 3b-d). Together, these findings identify that HIF stabilisation is required to orchestrate induction of *PLOD2* and *LOXL2* gene expression in human lung fibroblasts.

**Figure 3.**
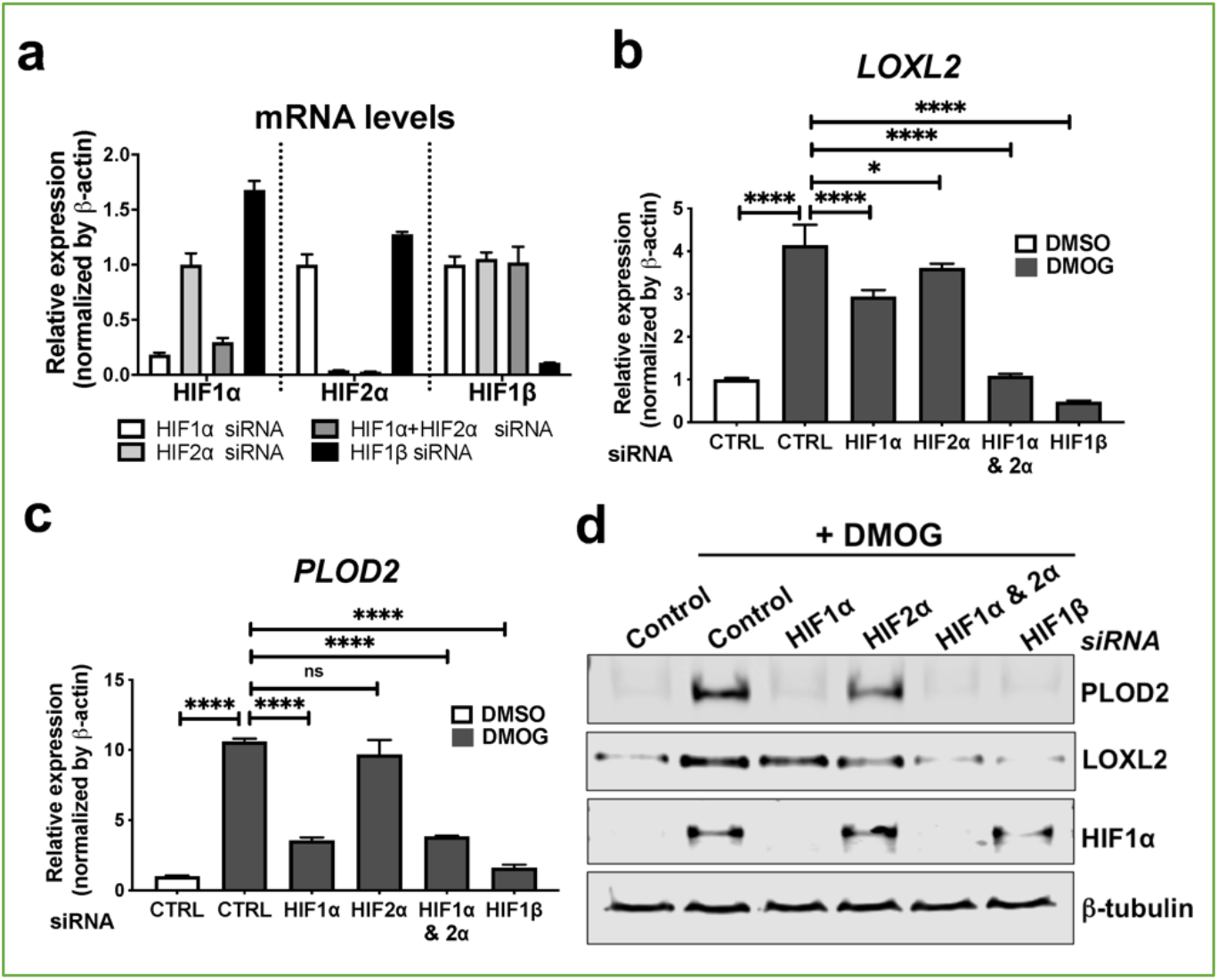
HIF pathway activation regulates PLOD2 and LOXL2 expression in lung fibroblasts from patients with IPF. (A) Fold change in mRNA levels of *HIF1α, HIF2α*, and *HIF1β* in primary human lung fibroblasts from patients with IPF transfected with indicated SiRNA followed by treatment with DMOG. β-actin-normalised mRNA levels in control cells were used to set the baseline value at unity. Data are mean ± s.d. n = 3 samples per group. (B, C) Fold change in mRNA levels of *LOXL2* (B) and *PLOD2* (C) in IPF fibroblasts transfected with indicated siRNA followed by treatment with DMOG or vehicle control. β-actin-normalised mRNA levels in control cells were used to set the baseline value at unity. Data are mean ± s.d. n = 3 samples per group. ns (not significant, p>0.05); *p<0.05; ****p<0.0001 by Dunnett’s multiple comparisons test. (D) Protein expression of PLOD2, LOXL2 and HIF1α and β-tubulin in IPF fibroblasts transfected with indicated siRNA followed by treatment of DMSO or DMOG. β-tubulin was used as a loading control. The full blots are shown in Figure 3-source data 1.

### HIF pathway activation and TGFβ1 synergistically increase PLOD2 expression

Given that TGFβ1 strongly induced major collagen fibrillogenesis genes whilst HIF pathways most strongly induced PLOD2 and LOXL2 expression, we investigated the effects of activating these pathways individually or in combination using lung fibroblasts from patients with IPF. The effect of DMOG or IOX2 in the presence or absence of TGFβ1 upon PLOD2 and LOXL2 induction was comparable to that identified using normal control lung fibroblasts (Fig. 4a-c). When combined, a synergistic effect upon the induction of PLOD2 expression was apparent which was greater than either pathway alone (Fig. 4a and b), whilst TGFβ1 alone was sufficient to induce interstitial collagen gene (*COL1A1*) expression (Supplementary Figure 3). HIF stabilisation significantly increased the ratio of *PLOD2* and *LOXL2* gene expression relative to fibrillar collagen (*COL1A1*) gene expression while TGFβ1 did not (Fig. 4d-e), suggesting that TGFβ activity alone may be insufficient to promote the altered collagen cross-linking that is present in IPF lung tissue. Together these findings identify that whilst TGFβ1 has a dominant role in increasing the rate of synthesis of major fibrillar collagens, HIF pathways may have a key role in regulating pathological post-translational modifications and collagen structure in lung fibrosis.

**Figure 4.**
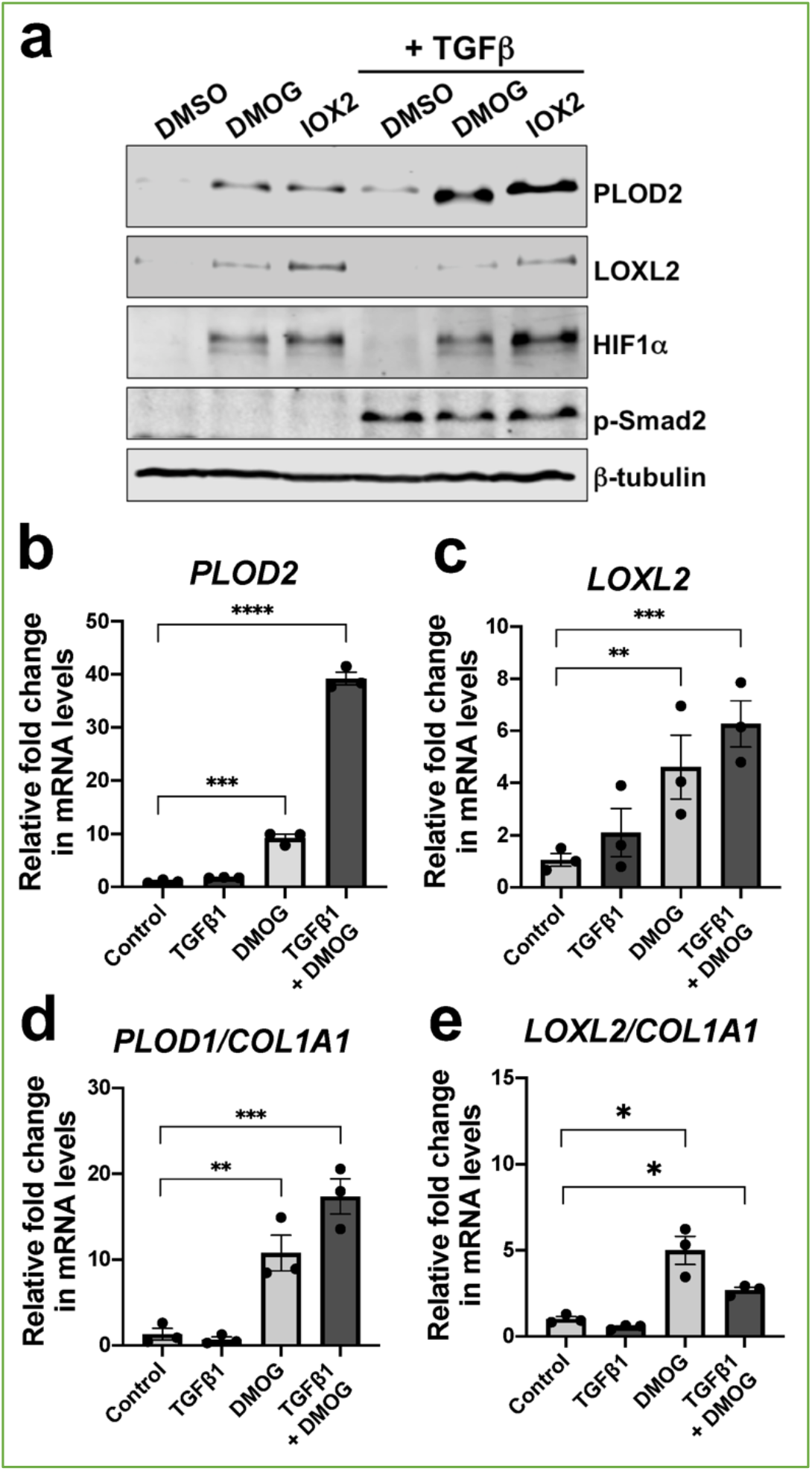
HIF pathway activation promotes increased pyridinoline post-translational gene modification expression relative to fibrillar collagen expression. (A) Protein expression of PLOD2, LOXL2 and HIF1α, phospho-Smad2 (p-Smad2) and β-tubulin in primary human lung fibroblasts from patients with IPF cultured in the presence of DMOG, IOX2, or vehicle control in the absence or presence of TGFβ1 for 48 hours. β-tubulin was used as a loading control. The full blots are shown in Figure 4-source data 1. (B-C) Relative gene expression of *PLOD2* and *LOXL2* was investigated in lung fibroblasts from IPF donors (n=3 across 2 independent experiments) exposed to 48 hours of media control, TGFβ1, DMOG or combined TGFβ1 and DMOG conditions using the ΔΔCt method. Bars indicate geometric means. Data are mean ± s.d. **p<0.01; ***p<0.001; ****p<0.0001 by Dunnett’s multiple comparisons test. (D-G) Expression of *PLOD2* and *LOXL2* from (B-C) was divided by *COL1A1* expression (shown in Supplementary Fig. 3) to calculate proportionate expression changes of cross-linking enzymes relative to collagen fibrillogenesis gene expression. Bars indicate geometric mean. Grouped analysis was performed using Dunnett’s multiple comparison test. ** p < 0.01, *** p < 0.001

### HIF pathway activation alters collagen structure-function and increases tissue stiffness

To investigate whether HIF pathway activation acts as a ‘second hit’ to drive pathologic collagen crosslinking by disproportionate induction of ‘bone-type’ collagen-modifying enzymes relative to TGFβ-induced collagen fibril synthesis, we employed our long-term 3D *in vitro* model of lung fibrosis using primary human lung fibroblasts from patients with IPF, so allowing direct evaluation of pyridinoline cross-linking, collagen nanostructure, and tissue biomechanics^1^. We employed the selective HIF-prolyl hydroxylase 2 inhibitor IOX2 to test within the *in vitro* fibrosis model, confirming HIF stabilisation by IOX2 following 2-week culture, and that in combination with TGFβ1 this promoted PLOD2 and LOXL2 expression (Supplementary Fig. 4). Following 6 weeks of culture with TGFβ1 in the absence (control) or presence of IOX2 to drive HIF pathway activation, mature pyridinoline cross links (DPD/PYD) were significantly increased by the addition of IOX2 (Fig. 5a) and these achieved a level comparable to our previous findings in IPF tissue^1^. The biomechanical consequence of HIF stabilisation by IOX2 treatment was then investigated with parallel plate compression testing, identifying a greater than 3-fold increase in tissue stiffness by the addition of IOX2 (Fig. 5b), with the mean (± SEM) compressive modulus measurement following IOX2 treatment of (107.1 ±10.7) kPa comparable to the maximal stiffness of between 50 and 150 kPa we and others have previously identified in highly fibrotic areas in IPF tissue^11^.

**Figure 5.**
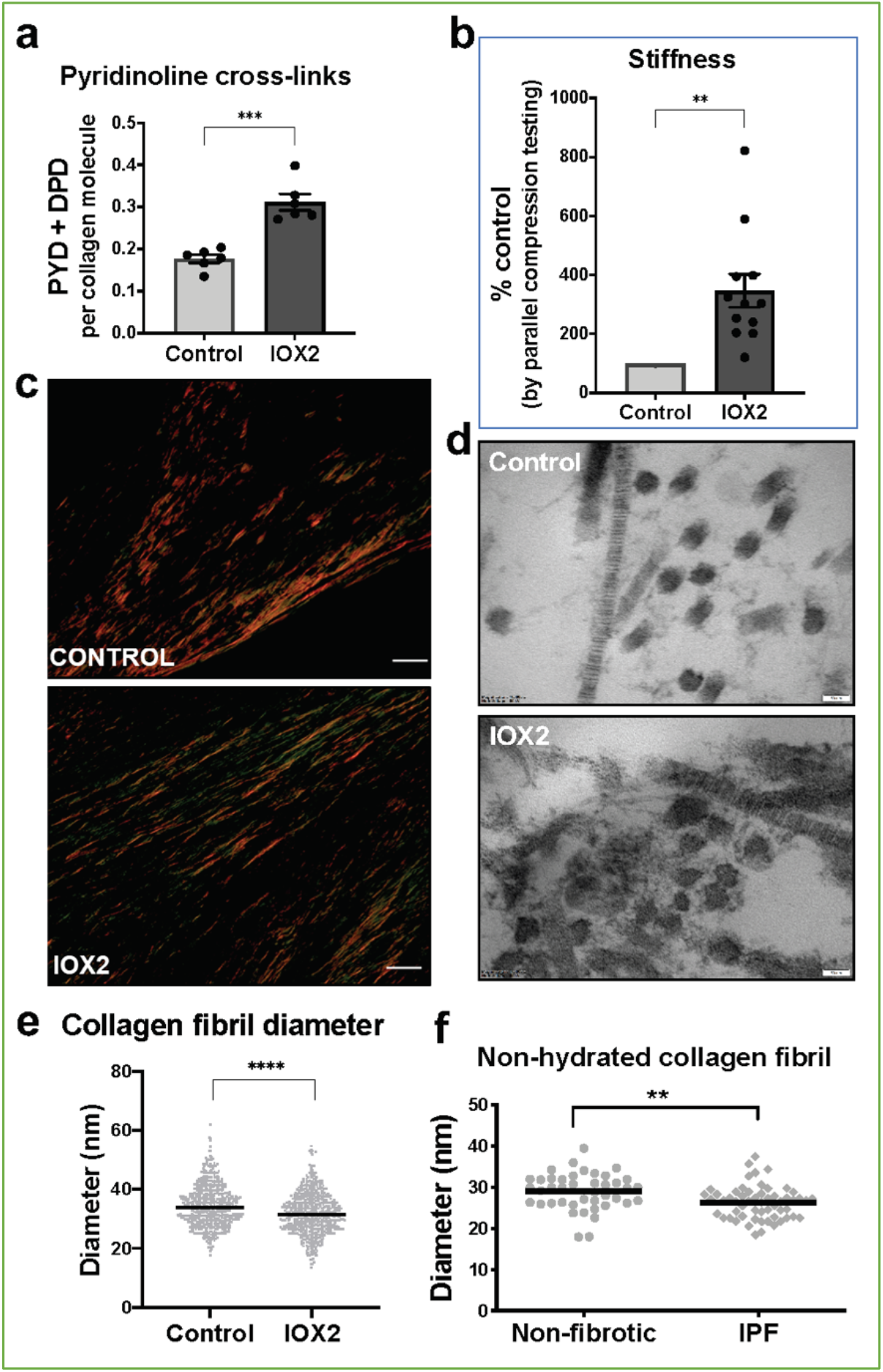
HIF pathway activation promotes pyridinoline cross-linking, alters collagen nano-architecture, and increases tissue stiffness. Lung fibroblasts from IPF patients (n = 3 donors, two experiments per donor) were used in the 3D model of fibrosis in the presence of IOX2 or vehicle control as indicated. Bars indicate geometric mean +s.e.m. Analysis was performed using a Mann-Whitney t-test (2-tailed) **p<0.01; ***p<0.001; ****p<0.0001. (A) Total mature trivalent (PYD + DPD) collagen cross-links determined by ELISA. n= 6 samples from 3 IPF donors. (B) Tissue stiffness measured from parallel-plate compression testing (n = 12 samples from 3 IPF donors) determined by Young’s modulus and represented as proportion of control. (C) Representative images of histological sections of samples stained with picrosirius red and imaged under polarised light. Scale bar 20 μm. (D) Representative electron microscopy images of collagen fibrils. Scale bar 50 nm. (E) Collagen fibril diameter measured in transverse section (300 fibrils for each condition from each of 2 IPF donors, measured by a blinded investigator). (F) Atomic force microscopy indentation modulus of collagen fibrils (3–7 fibrils per donor) from control (n = 42 fibrils from eight donors) or IPF lung tissue (n = 57 fibrils from 10 donors) under non-hydrated conditions; each data point represents the mean of 30 to 50 force-displacement curves per fibril.

We next assessed collagen morphology. When visualised by polarised light Picrosirius red microscopy (Fig. 5c), highly organised collagen fibrils were evident in vehicle-treated fibrotic control cultures as well as in those treated with IOX2 with no apparent morphological differences. In contrast ultrastructural analysis of the collagen fibrils using electron microscopy identified a change in collagen nanostucture with a significant decrease in fibril diameter (Fig. 5d and e) when pyridinoline cross-linking was increased by IOX2, consistent with our previous observation that fibril diameter is increased by inhibition of pyridinoline cross-linking^1^. In support of the disease relevance of our *in vitro* findings, non-hydrated collagen fibrils from patients with IPF have reduced diameters when measured by atomic force microscopy (Fig. 5f), consistent with our previous findings that hydrated collagen fibrils extracted from IPF lung tissue have a reduced diameter compared to control samples^1^. Together, these data identify HIF pathway activation to be a key regulator of pyridinoline cross-link density, collagen fibril nano-architecture, and tissue stiffness.

### Pseudohypoxia and loss of FIH activity promotes HIF pathway activation in lung fibroblasts

Whilst canonical HIF pathway activation was observed in lung fibroblasts under hypoxic conditions, elevated levels of HIF1*α* and HIF2*α* in IPF fibroblasts under normoxic conditions have recently been reported^12^, suggesting a pseudohypoxic state i.e. a state in which cells express hypoxia-associated genes and proteins, regardless of the oxygen status^13^. To further investigate this possibility, we employed gene set variation analysis (GSVA) using a validated 15-gene HIF/hypoxia gene expression signature^14^ to published datasets, identifying that cultured fibroblasts from patients with a usual interstitial pneumonia pattern of fibrosis or systemic sclerosis associated lung fibrosis have a significantly increased HIF score compared to cultured control fibroblasts (Fig. 6a), consistent with an oxygen independent increase in HIF activity. Furthermore, there was a significant increase in the HIF score in lung mesenchymal stromal cells of patients with progressive lung fibrosis compared to those with stable fibrosis (Fig. 6b), suggesting that HIF pathway activation may be required for fibrosis progression.

**Figure 6.**
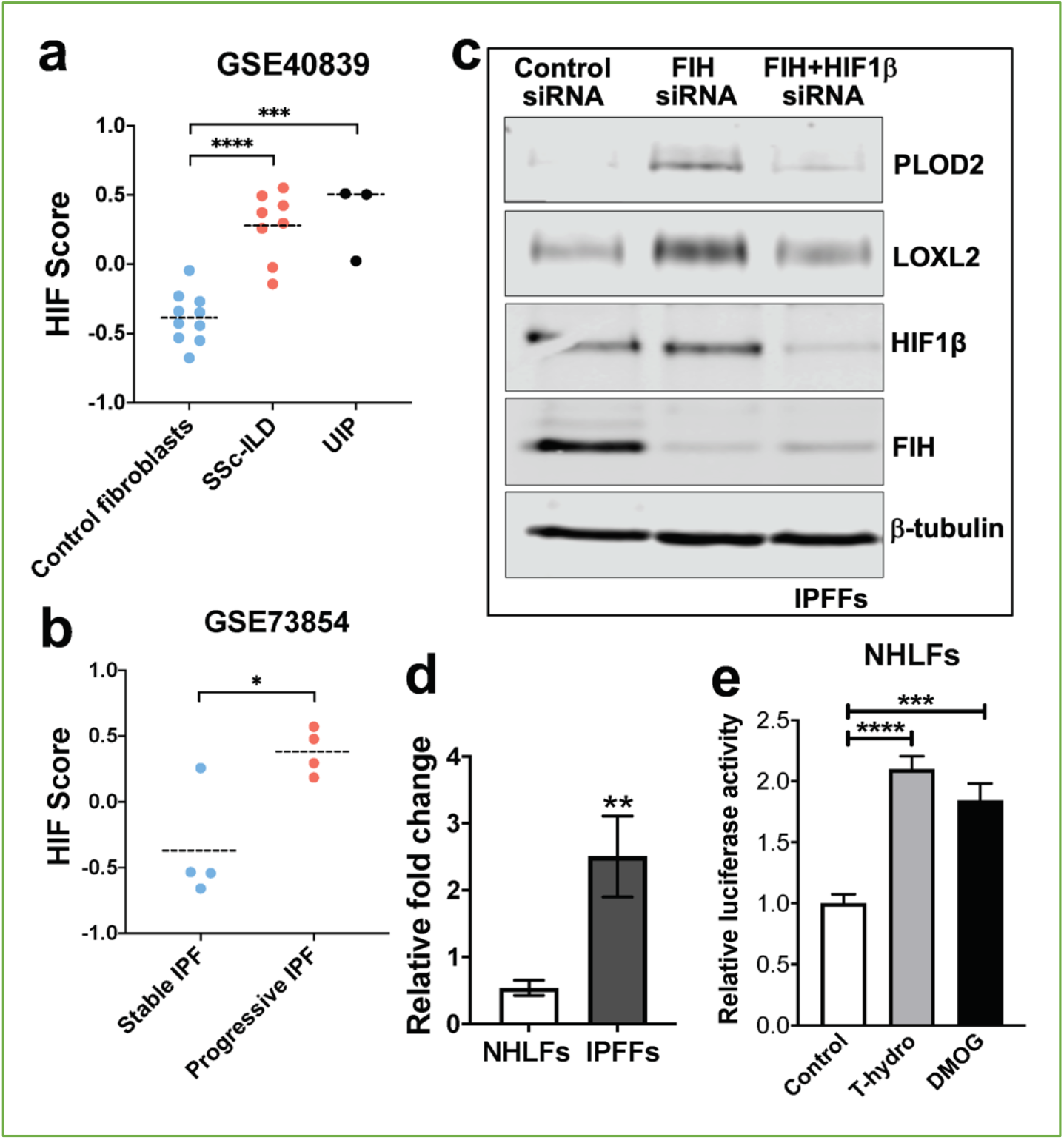
Pseudohypoxia and loss of FIH activity promotes HIF pathway signalling in IPF fibroblasts. (A) HIF GSVA scores calculated in human lung fibroblasts derived from control or patients with interstitial lung disease (scleroderma lung or a usual interstitial pneumonia / IPF pattern) (GSE40839). Data are mean ± s.d. ***p<0.001; ****p<0.0001 by Dunnett’s multiple comparisons test. (B) HIF GSVA scores calculated in human bronchoalveolar lavage derived mesenchymal stromal cells from patients with stable and progressive IPF (GSE73854). Data are mean ± s.d. *p<0.05 by unpaired t test. (C) Protein expression of PLOD2, LOXL2, HIF1β, FIH and β-tubulin in lung fibroblasts from patients with IPF transfected with indicated siRNA. β-tubulin was used as a loading control. The full blots are shown in Figure 6-source data 1. (D) FIH reporter assays in NHLFs or IPFFs. Values represent relative fold of firefly luciferase in relation to Renilla luciferase, normalised against control (1.0). Data are mean ± s.d. n = 3 samples per group. **p< 0.01 by unpaired t test. (E) FIH reporter assays in NHLFs with indicated treatment (hydrogen peroxide, DMOG, or vehicle control). Values represent relative fold of firefly luciferase in relation to *Renilla* luciferase, normalised against control (1.0). Data are mean ± s.d. n = 3 samples per group. ***p<0.001; ****p<0.0001by Dunnett’s multiple comparisons test.

To examine the mechanism underlying pseudohypoxic HIF activity in lung fibrosis, we investigated the role of Factor Inhibiting HIF (FIH), a Fe (II)- and 2-oxoglutarate (2-OG)-dependent dioxygenase, which regulates HIF activity via hydroxylating a conserved asparagine (Asn) residue within the HIFα C-terminal activation domain (CAD), a post-translational modification that blocks interactions between the HIFα-CAD and CBP/p300^15-19^. Whilst oxygen tension is the classical regulator of FIH activity, oxidative stress can inactivate FIH so promoting HIF activity under normoxic conditions^20^.

Initially, to investigate the potential role of reduced FIH activity in regulating collagen post-translational modifications, we silenced FIH under normoxic conditions, identifying that loss of FIH was sufficient to induce both PLOD2 and LOXL2 expression, and that this effect required HIF activity, since HIF1β knockdown prevented their induction (Fig. 6c). Whilst FIH is stably constitutively expressed across tissues^21, 22^, the activity levels of FIH can vary^23-25^; thus we compared FIH activity in control or IPF fibroblasts using a UAS-luc/GAL4DBD-HIF1αCAD binary reporter system^26^. In this assay, the activity of FIH is monitored by a Gal4-driven luciferase reporter that registers the activity of the heterologous Gal4-HIF-CAD fusion protein. Inhibition of FIH leads to a reduction in hydroxylation at Asn-803 of the HIF-CAD fusion, which permits increased recruitment of the transcriptional co-activators p300/CBP and enhanced reporter gene activity (Supplementary Figure 5). Consistent with a loss of function of FIH in lung fibrosis, we found FIH activity was significantly reduced in fibroblasts from patients with IPF compared to control fibroblasts (Fig. 6d). We further confirmed that a reduction in FIH activity in normal lung fibroblasts could be caused under normoxia by oxidative stress, achieving a level of HIF activation comparable to treatment with the hypoxia mimetic DMOG (Fig. 6e). Thus, in lung fibroblasts a reduction in FIH activity may promote HIF pathway activation to dysregulate collagen structure-function.

### HIF pathway activation localises in areas of active fibrogenesis to cells co-expressing LOXL2 and PLOD2

To support our in vitro studies, we investigated for evidence that HIF regulates *PLOD2* and *LOXL2* expression within the fibroblast foci of human IPF lung tissue. To assess for HIF activity we applied GSVA using the 15-gene HIF/hypoxia gene expression signature^14^ to the transcriptome of each fibroblast focus, identifying that the HIF signature score, but not TGFβ score, significantly correlated with *LOXL2*/*PLOD2* expression (Fig. 7a and b). Furthermore, analysis of serial tissue sections using immunohistochemistry identified that HIF1*α* and the HIF pathway activation marker gene carbonic anhydrase IX (CA-IX) were expressed within fibroblast foci^7, 27^, and that this expression localised to cells co-expressing *LOXL2* and *PLOD2* mRNA (Fig. 7c). Finally, as FIH is more sensitive to inhibition by oxidative stress^20^ whilst PHDs are more sensitive to hypoxia we investigated whether HIF activation occurs in lung mesenchymal cells in the context of oxidative stress. We applied GSVA to a published single cell RNAseq dataset (114,396 cells) from 10 control and 20 fibrotic lungs which identified 31 cell types including 4 fibroblast types (fibroblasts, myofibroblasts, Hyaluronan Synthase 1 (HAS1) high and Perilipin 2 (PLIN2)+ fibroblasts) (Supplementary Fig. 6)^28^. Applying HIF signature or upregulated oxidative stress gene expression signatures to this dataset, we identified that, compared to fibroblasts and myofibroblasts, the HAS1 high and PLIN2+ cells, whose presence was almost exclusively derived from the IPF lung tissue, had significantly increased HIF and upregulated oxidative stress scores (Fig. 8a-d) and that these significantly correlated (Fig. 8e), consistent with an increase in pseudohypoxic HIF activity in these disease-specific mesenchymal cell types.

**Figure 7.**
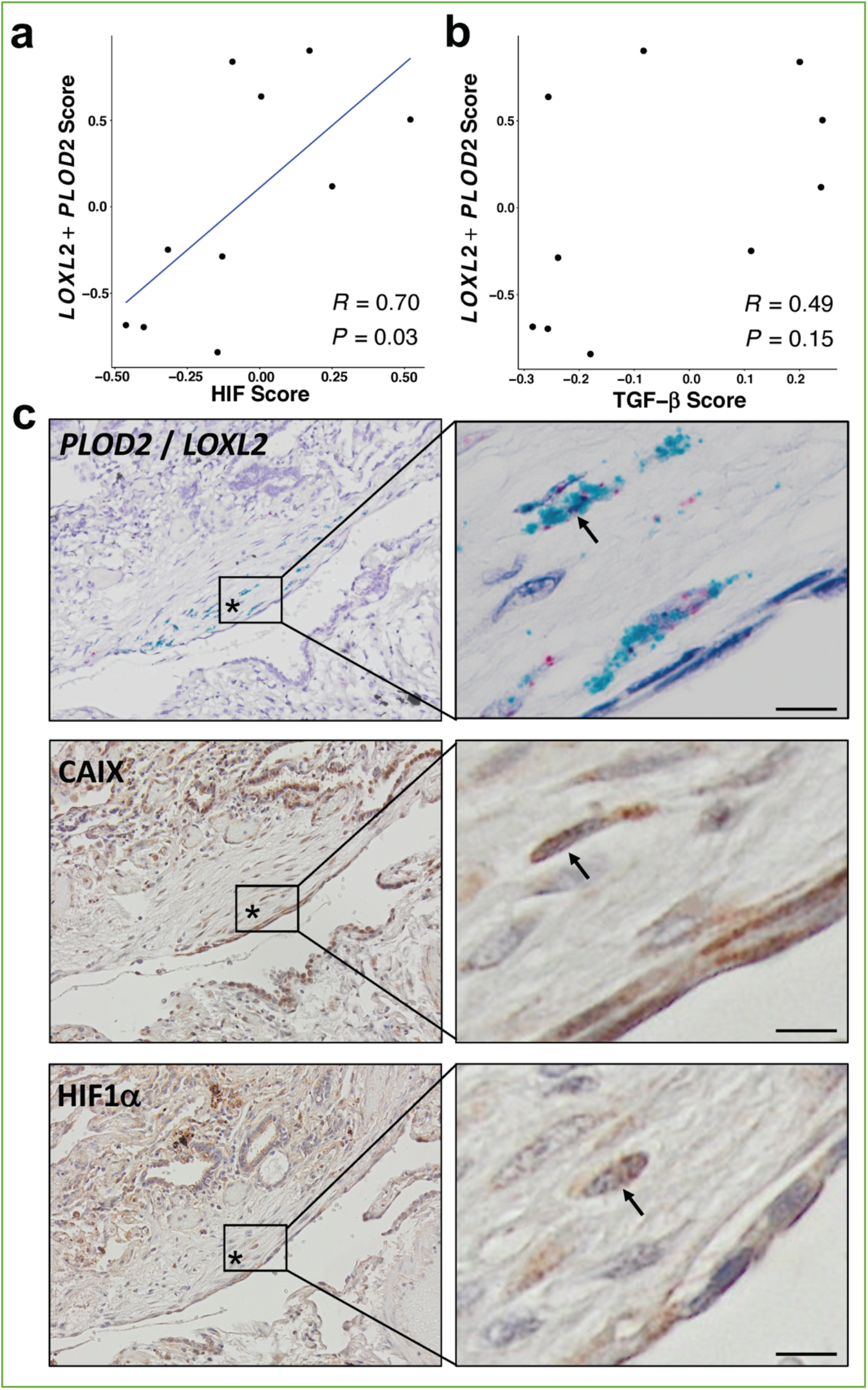
HIF pathway activation localises in areas of active fibrogenesis to cells co-expressing LOXL2 and PLOD2. (A-B) Scatterplots showing correlations between *LOXL2*/*PLOD2* expressions and HIF scores (A) or TGF*β* scores (B) in IPF fibroblast foci (n= 10) using Spearman correlation coefficient. (C) Representative images of serial sections of lung tissue from patients with IPF (n=3). mRNA expression of *PLOD2* (red chromagen) and *LOXL2* (green chromagen) using RNAscope^®^ RNA in-situ hybridisation with immunohistochemical staining for Carbonic anhydrase IX (CA-IX) and HIF1α using DAB (brown). A fibroblastic focus is identified by *. Scale bar 20 μm.

**Figure 8:**
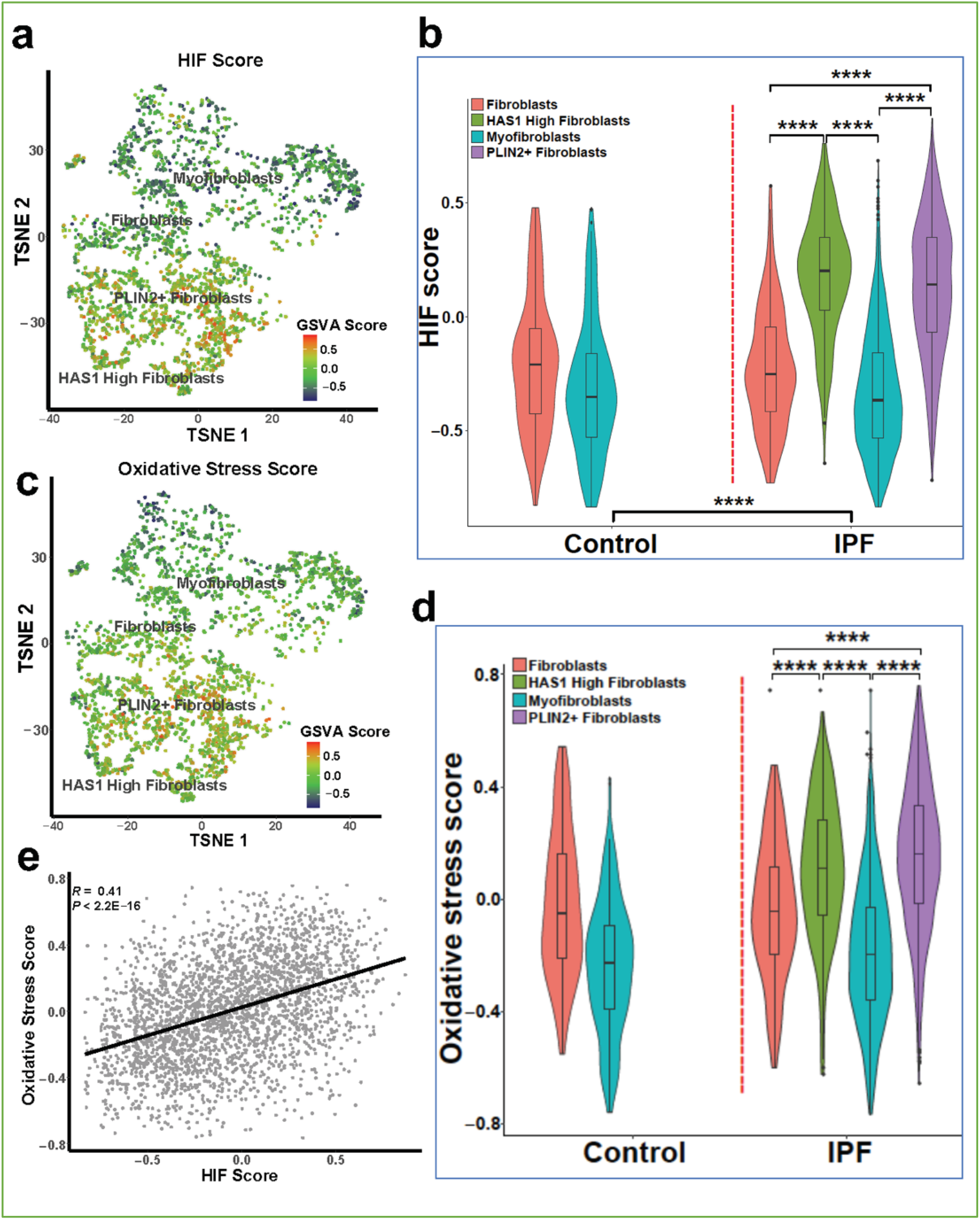
Gene set variance analysis of single-cell RNAseq fibroblast populations identifies co-enrichment of HIF score and oxidative stress genes. A: HIF score GSVA in control and IPF fibroblasts sequenced by single-cell RNAseq (GSE135893). Colours correspond to calculated GSVA score for each cell. B: Plot of mean HIF GSVA scores for each fibroblast type in control and IPF fibroblast cell populations and compared using Dunnett’s multiple comparison test, ****P < 0.0001. C: GSVA scores for genes upregulated in IPF in this dataset associated with the Gene Set: HALLMARK_REACTIVE_OXYGEN_SPECIES_PATHWAY (M5938). D: Plot of upregulated oxidative stress GSVA scores for each fibroblast type in control and IPF cells. E: Correlation plot of HIF score vs upregulated oxidative stress GSVA score for single cell RNAseq data. Correlation coefficient is Pearson’s product-moment coefficient.

## Discussion

We previously reported that altered collagen fibril nanoarchitecture is a core determinant of dysregulated ECM structure-function in human lung fibrosis^1^. Here, through ex vivo models, bioinformatics and human lung fibrosis tissue studies, we extend these observations to identify that HIF pathway activation promotes pathologic collagen crosslinking and tissue stiffness by disproportionate induction of ‘bone-type’ collagen-modifying enzymes relative to TGFβ-induced collagen fibril synthesis. Furthermore, this may occur via pseudohypoxic oxygen-independent mechanisms involving loss of FIH activity that can occur due to oxidative stress, which is thought to play a significant role in IPF pathogenesis^29^. Consistent with this, oxidative stress is increased in subpopulations of IPF fibroblasts whilst FIH activity is significantly reduced in fibroblasts from patients with lung fibrosis resulting in HIF activation under normoxic conditions. Thus, we provide evidence that dysregulated HIF activity is a core regulator of ECM structure-function in human lung fibrosis, and that this may be a key determinant of pathologic tissue stiffness and progressive human lung fibrosis.

TGFβ is a multifunctional growth factor with key roles in normal development and wound healing. It is also considered the prototypic profibrogenic cytokine that promotes increased ECM deposition and has been associated with fibrosis across multiple organs^6^. We identified that in lung fibroblasts, TGFβ1 increased fibrillar collagen mRNA transcription but proportionally its effects on *PLOD2* or *LOXL2* were more limited, suggesting that TGFβ pathway activation alone may be insufficient to cause a substantial increase in bone-type pathologic collagen crosslinking. Rather, HIF pathway activation was required to disproportionately induce PLOD2/LOXL2 relative to interstitial collagen fibril synthesis, so promoting ‘bone’ type collagen fibrillogenesis, altering collagen fibril nanostructure, and increasing tissue stiffness. While TGFβ has been reported to cause HIF stabilization^30^, our findings suggest that this effect is modest and that further HIF activation is required to drive matrix stiffening. This would be in keeping with a recent observation of an hierarchical relationship in which HIF proteins play a more powerful role in the induction of PLOD2 expression, with HIF transcription factors dominating the effect of TGFβ stimulated SMAD proteins^31^. Thus, we propose that HIF pathway activation acts as a key pathologic ‘second hit’ which disrupts the normal wound healing role of TGFβ by altering collagen fibril nanoarchitecture so dysregulating ECM structure-function and promoting progressive lung fibrosis. In keeping with this concept, GSVA using a validated HIF score^14^ applied to microarray data for lung mesenchymal stromal cells showed that HIF activity was increased in cells from patients with progressive lung fibrosis compared with those with stable disease.

We investigated the functional consequences of our findings by employing our long-term 3D *in vitro* model of lung fibrosis, identifying that HIF pathway activation increased pyridinoline cross-links to a level comparable to that identified in IPF tissue, and that this was associated with an increase in tissue stiffness comparable to the extremes of stiffness identified in IPF tissue together with a reduction in fibril diameter similar to those present in IPF lung tissue. Together these observations support the human disease relevance of HIF pathway activation to IPF and define conditions for future mechanistic studies whereby the 3D *in vitro* model recapitulates key features of dysregulated collagen structure-function in IPF. Hypoxia has been proposed to have a pathogenetic role in lung fibrosis through mechanisms including fibroblast proliferation, augmented ER stress, epithelial-mesenchymal transition, and glycolytic reprogramming^7, 32-34^. Furthermore, a number of reports have proposed that cross-talk between TGFβ and hypoxia may promote fibrosis, with hypoxia and TGFβ1 synergistically increasing myofibroblast marker expression^35^, promoting experimental nickel oxide nanoparticle-induced lung fibrosis^36^, and HIF1α mediating TGF-β-induced PAI-1 production in alveolar macrophages in the bleomycin model of lung fibrosis^37^.

The HIF signalling pathway has been reported to be active in lungs and fibroblasts from IPF patients, as determined by the abundance of HIF alpha subunits 1 and 2^7^. These findings are consistent with our own observations of increased expression of the HIF-responsive gene, CA-IX. Here we extended these observations by showing that in lung fibrosis, loss of FIH activity either by siRNA-mediated knockdown or exposure to oxidative stress induces HIF pathway activation independently of oxygen tension, so dysregulating collagen fibrillogenesis under normoxic conditions. FIH negatively regulates HIF activity by hydroxylation of N803, preventing the interaction of the HIFα CAD with CBP/p300^15-19^. Whilst oxygen tension is the classical regulator of FIH activity, oxidative stress can inactivate FIH so promoting HIF activity^20^. Oxidative stress has been implicated as an important profibrotic mechanism in the lungs and other organs^29, 38, 39^. It can arise from exposure to environmental toxins (e.g. air pollution, tobacco, asbestos, silica, radiation, and drugs such as bleomycin) or from endogenous sources including mitochondria, NADPH oxidase (NOX) activity, and/or inadequate or deficient antioxidant defenses^29^. In our own bioinformatic studies, we observed subsets of disease-specific fibroblasts with elevated scores for oxidative stress and these same populations had evidence of HIF pathway activation.

To our knowledge whether perturbations in FIH activity could contribute to fibrosis has not been investigated previously. Whilst our studies have focussed upon HIF pathways and collagen, functionally FIH, via both HIF-dependent and HIF-independent pathways, has been reported to regulate metabolism^40-43^, keratinocyte differentiation^44^, vascular endothelial cell survival^45^, tumour growth^46, 47^ and metastasis^48^ as well as Wnt signalling^49^, suggesting that the loss of FIH activity we have identified could have pleiotropic effects in lung fibrosis which merits further investigation.

In summary, this study identifies that HIF pathway activation via oxygen dependent and oxygen independent mechanisms promotes ‘bone type’ collagen nano-architecture which is a defining feature of human lung fibrosis that dysregulates ECM structure-function to promote progressive lung fibrosis. Our findings suggest that therapeutically targeting HIF pathway activation might restore ECM homeostasis and so prevent fibrosis progression.

## MATERIALS AND METHODS

### Lung Tissue Sampling

Human lung experiments were approved by the Southampton and South West Hampshire and the Mid and South Buckinghamshire Local Research Ethics Committees (ref 07/H0607/73), and all subjects gave written informed consent. Clinically indicated IPF lung biopsy tissue samples deemed surplus to clinical diagnostic requirements were formalin fixed and paraffin embedded. All IPF samples were from patients subsequently receiving a multidisciplinary diagnosis of IPF according to international consensus guidelines.

### Transcriptomic analysis of in situ IPF fibroblast foci

We analysed a transcriptomic data set that we have recently established (GSE169500). Briefly, laser capture microdissection was performed upon Formalin-Fixed Paraffin-Embedded (FFPE) control non-fibrotic lung tissue (alveolar septae, (n=10)) and usual interstitial pneumonia/idiopathic pulmonary fibrosis FFPE lung tissue (fibroblast foci, (n=10) and adjacent non-affected alveolar septae, (n=10)). Total RNA was isolated, cDNA libraries were prepared using Ion Ampli-Seq-transcriptome human gene expression kit (Life Technologies, Paisley, UK) and sequenced using Ion Torrent Proton Sequencer. A two-stage mapping strategy was used to map the reads to UCSC hg19 human genome. Cufflinks was used to calculate Fragments per Kilobase of exon per Million (FPKM) values.

### RNA in-situ hybridisation

Simultaneous in situ detection of the LOXL2 and PLOD2 mRNA on human IPF formalin fixed paraffin embedded tissue sections were performed using duplex RNAscope© technology (Advanced Cell Diagnostics, Biotechne, Abingdon, UK). *LOXL2* was detected by C1-probe (Probe-Hs-LOXL2-C1, 311341) and *PLOD2* was detected by C2-probe (Probe-Hs-PLOD2-C2, 547761-C2). Briefly, 5-μm human IPF lung tissue sections were baked at 60C, deparaffinized in xylene, followed by dehydration in an ethanol series. Target retrieval, hybridization with target probes, amplification, and chromogenic detection were performed according to the manufacturer’s recommendations (RNAscope 2.5 Duplex Detection protocol for FFPE tissues). Sections were counterstained with Gill’s Hematoxylin, and mounted with Vectamount permanent mounting medium prior to imaging. Assays were performed with duplex positive (*PPIB* and *POLR2A*) and negative controls. Images were acquired using an Olympus Dotslide Scanner VS110 (Olympus UK, Southend-on-Sea, UK).

### 2D Cell culture, reagents and transfections

Primary fibroblast cultures were established from lung parenchyma tissue of patients with IPF obtained by video-assisted thoracoscopic lung biopsy at University Hospital Southampton or non-fibrotic control lung parenchyma tissue (macroscopically normal lung sampled remote from a cancer site in patients undergoing surgery for early stage lung cancer)^1, 3, 50, 51^. MRC5 lung fibroblasts were obtained from the European Collection of Authenticated Cell Cultures (ECACC). All cultures were tested and free of mycoplasma contamination. Fibroblasts were cultured in Dulbecco’s Modified Eagle’s Medium (DMEM) supplemented with 10% foetal bovine serum (FBS), 50 units/ml penicillin, 50μg/ml streptomycin, 2mM L-glutamine, 1mM sodium pyruvate and 1x non-essential amino acids (DMEM/FBS) (Life Technologies, Paisley, UK). All cells were kept at 37 °C and 5% CO2. Hypoxic incubation of cells was carried out in a H35 Hypoxystation (Don Whitley Scientific) in which cells were cultured in humidified atmosphere of 1% O2, 5% CO2, and 94% N2 at 37 °C. Following hypoxic incubation, cells were kept in hypoxic condition until samples were collected.

For pro-fibrogenic mediator studies control lung fibroblasts were treated in the presence of EGF (R&D systems, 236-GMP-200, 10ng/mL), TGFβ1 (R&D systems, 240-GMP-010, 10ng/mL), Dimethyloxaloylglycine (DMOG) (Merck, CAS89464-63-1, 1mM), Wnt3a (R&D systems, 5036-WN-010, 100ng/mL), Wnt5a (R&D systems, 645-WN-010, 100ng/mL), or vehicle control (DMSO). For subsequent HIF studies fibroblasts were treated in the presence of DMOG (1mM), IOX2 (50 μM or 250 μM) or vehicle control (DMSO)

Short interfering RNA (siRNA) oligos against HIF1A (HIF1α) (MU-00401805-01-0002), EPAS1 (HIF2α) (MU-004814-01-0002), ARNT (HIF1β) (MU-007207-01-0002) and HIF1AN (FIH) (MU-004073-02-0002), LOXL2(L-008020-01-0005) were purchased from Dharmacon, Cambridge, UK. Sequences are available from Dharmacon, or on request. As a negative control, we used siGENOME RISC-Free siRNA (Dharmacon, D-001220–01). Human lung fibroblasts were transfected with the indicated siRNA at a final concentration of 35 nM using Lipofectamine RNAiMAX reagent (Invitrogen).

### Reporter assay

FIH activity was evaluated using a UAS-luc/GAL4DBD-HIF1αCAD binary reporter system^26^. For the luciferase reporter assays in a 24 well plate, human lung fibroblasts (NHLFS or IPFFs) were reverse transfected using Lipofectamine 3000 (Invitrogen) with 50 ng of phRL-CMV (Promega UK, Southampton, UK), which constitutively expresses the *Renilla* luciferase reporter, plus 225 ng of plasmid-GAL4DBD-HIF1αCAD. and 225 ng of plasmid-UAS-luc per well. After 24-hour of transfections, a final concentration of 1mM of DMOG, 1mM DMSO or 20μM freshly prepared T-hydro (tert-butyl hydroperoxide) (Sigma-Aldrich, Poole, UK) was dosed for 16 hours. T-hydro was added to the cells every 2 hours. Finally, the transcriptional assay was carried out using the Dual-Luciferase reporter assay system (Promega) following the manufacturer’s protocol.

### HIF score, TGFβ score and Oxidative Stress GSVA Analyses

Raw CEL files for GSE73854 and GSE40839 were downloaded from GEO and imported into RStudio (version 3.6). Raw data were normalised by Robust Multi-array Average (RMA) function in the affy package (version 1.64.0). Multiple probes relating to the same gene were deleted and summarised as the median value for further analysis.

A 15-gene expression signature (*ACOT7, ADM, ALDOA, CDKN3, ENO1, LDHA, MIF, MRPS17, NDRG1, P4HA1, PGAM1, SLC2A1, TPI1, TUBB6* and *VEGFA*) was selected to classify HIF activity^14^. All parameters and variables can be found in the accompanying file (Source code 1). This gene signature was defined based on knowledge of gene function and analysis of in vivo co-expression patterns and was highly enriched for HIF-regulated pathways. The HIF score for each sample was calculated by using gene set variation analysis (GSVA)^52^ based on this 15-gene expression signature. The TGF*β* score for each sample was calculated by using GSVA based on a list of gene from Gene Set: HALLMARK_TGF_BETA_SIGNALING (M5896). All parameters and variables can be found in the accompanying file (Source code 2). The Student t-test was used to evaluate the statistical difference in HIF scores between different conditions.

For single cell transcriptomic analyses raw CEL files for GSE135893 were downloaded from GEO. Data was processed using the Seurat R package (v3.2.1) in R version 4.0.2. Cell types were assigned based on the published metadata^28^. Fibroblast counts data were log-normalised, variable genes quantified and principal component analysis performed on these variable genes. T-stochastic nearest neighbour embedding (t-SNE) dimensional reduction was performed on the top 15 principal components to obtain embeddings for individual cells. GSVA was performed using the 15 genes used for HIF score calculation as above. An oxidative stress score for each cell was calculated using GSVA based on a list of genes upregulated in IPF cell populations (*ABCC1, CDKN2D, FES, GCLC, GCLM, GLRX2, HHEX, IPCEF1, JUNB, LAMTOR5, LSP1, MBP, MGST1, MPO, NDUFA6, PFKP, PRDX1, PRDX2, PRDX4, PRNP, SBNO2, SCAF4, SOD1, SOD2, RXN1, TXN, TXNRD1*) from Gene Set: HALLMARK_REACTIVE_OXYGEN_SPECIES_PATHWAY (M5938). All parameters and variables can be found in the accompanying file (Source code 3). Upregulated oxidative stress genes were those whose expression was higher in IPF populations than control. Calculated GSVA scores were mapped onto t-SNE plots. Student’s t-test was used to calculate statistical differences between GSVA scores of the different cellular populations.

### 3D in vitro model of fibrosis

Culture was performed as previously described^1^. Briefly, peripheral lung fibroblasts were obtained as outgrowths from surgical lung biopsy tissue of patients (n = 3 donors) who were subsequently confirmed with a diagnosis of IPF. All primary cultures were tested and free of mycoplasma contamination. The fibroblasts were seeded in Transwell inserts in DMEM containing 10% FBS. After 24 hr, the media was replaced with DMEM/F12 containing 5% FBS, 10 μg/ml L-ascorbic acid-2-phosphate, 10 ng/ml EGF, and 0.5 μg/ml hydrocortisone with or without 50 μM or 250 μM IOX2, as indicated; each experiment included a vehicle control (0.2% DMSO). TGF-β1 (3 ng/mL) was added to the cultures, and the medium replenished three times per week. After 2 weeks spheroids were lysed for western blotting. After 6 weeks the spheroids were either snap frozen for parallel-plate compression testing, analysis of cross-linking, and histochemical staining, or fixed using 4% paraformaldehyde for histochemistry or 3% glutaraldehyde in 0.1 M cacodylate buffer at pH 7.4 for electron microscopy.

### Reverse transcription quantitative polymerase chain reaction (RTqPCR)

RTqPCR was performed as previously described^3, 50, 51^. Primers and TaqMan probe sets were obtained from Primer Design, Southampton, UK (*LOXL2, COL1A1, Col3A1, PLOD2*), ThermoFisher Scientific, Reading, UK (*HIF1A, EPAS1, HIF1β*), and Qiagen, Manchester, UK (QuantiTect Primer Assays, *HIF1A, EPAS1, ARNT, LOXL2, PLOD2, CA9, ACTB*).

### Western blotting

Fibroblasts were lysed using 2x Laemmli SDS sample buffer or urea buffer (8 M Urea, 1 M Thiourea, 0.5% CHAPS, 50 mM DTT, and 24 mM Spermine). Western blotting of cellular lysates was performed for β-actin (1:100.000, Sigma-Aldrich, Poole, UK), LOXL2 (1:1000, R&D Systems, Abingdon, UK), HIF1α (1:1000, BD Biosciences, Wokingham, UK), FIH (1:200, mouse monoclonal 162C)^53^, β-tubulin (1:1000, Cell Signaling Technology, London, UK), HIF1 β (1:1000, Cell Signaling Technology), p-Smad2/3 (1:1000, Cell Signaling Technology), p-ERK (1:1000, Cell Signaling Technology), active β-catenin (1:1000, Cell Signaling Technology). Immunodetected proteins were identified using the enhanced chemiluminescence system (Clarity Western Blotting ECL Substrate, Bio-Rad Laboratories Ltd, Watford, UK) or Odyssey imaging system (LI-COR), and evaluated by ImageJ 1.42q software (National Institutes of Health).

### Immunofluorescence staining

Cells were fixed with 4% paraformaldehyde followed by permeabilization and staining with primary antibodies for LOXL2 (1:500, R&D Systems) and Tetramethylrhodamine (TRITC)-conjugated Phalloidin (1:1000, Millipore UK Limited, Watford, UK). The secondary antibodies used were Alexafluor 488 and 647 (1:1000, BioLegend UK Ltd, London, UK). Cell nuclei were counterstained with 4’,6-Diamidino-2-Phenylindole, Dihydrochloride (DAPI) (1:1000, Millipore UK Limited, Watford, UK). Cells were imaged using an inverted confocal microscope (Leica TCS-SP5 Confocal Microscope, Leica Microsystems).

### Immunohistochemistry

Control or IPF lung tissues (n = 3 donors) were fixed and embedded in paraffin wax; tissue sections (4μm) were processed and stained as previously described^3, 51^. Briefly, the tissue sections were de-waxed, rehydrated and incubated with 3% hydrogen peroxide in methanol for 10 min to block endogenous peroxidase activity. Sections were then blocked with normal goat serum and incubated at room temperature with a primary antibody against CA-IX (1:500, Novus Biologicals, Cambridge, UK) or HIF1*α* (1:500, Cayman Chemical, Michigan, USA), followed by a biotinylated secondary antibody (1:500, Vector Laboratories Ltd., UK); antibody binding was detected using streptavidin-conjugated horse-radish peroxidase and visualised using DAB before counter-staining with Gill’s Haematoxylin. Images were acquired using an Olympus Dotslide Scanner VS110.

### Protein, hydroxyproline and collagen cross-link assays

performed as previously described^1^.

### Parallel plate compression testing

performed as previously described^1^.

### Transmission electron microscopy

performed as previously described^1^.

### Atomic force microscopy nanoindentation imaging of individual non-hydrated collagen fibrils

performed as previously described^1^.

### Statistics

Statistical analyses were performed in GraphPad Prism v7.02 (GraphPad Software Inc., San Diego, CA) unless otherwise indicated. No data were excluded from the studies and for all experiments, all attempts at replication were successful. For each experiment, sample size reflects the number of independent biological replicates and is provided in the figure legend. Normality of distribution was assessed using the D’Agostino-Pearson normality test. Statistical analyses of single comparisons of two groups utilised Student’s t-test or Mann-Whitney U-test for parametric and non-parametric data respectively. Where appropriate, individual t-test results were corrected for multiple comparisons using the Holm-Sidak method. For multiple comparisons, one-way analysis of variance (ANOVA) with Dunnett’s multiple comparison test or Kruskal-Wallis analysis with Dunn’s multiple comparison test were used for parametric and non-parametric data, respectively. Results were considered significant if p<0.05, where *p<0.05, **p<0.01, ***p<0.001, ****p<0.0001.

## Acknowledgements

This project was supported by Medical Research Council (MR/S025480/1), the Wellcome Trust (100638/Z/12/Z), an Academy of Medical Sciences/the Wellcome Trust Springboard Award [SBF002\1038], and the AAIR Charity. CJB and LSND acknowledge the support of the NIHR Southampton Biomedical Research Centre. LY was supported by China Scholarship Council. YZ was supported by an Institute for Life Sciences PhD Studentship. FC was supported by Medical Research Foundation [MRF-091-0003-RG-CONFO]. ML was supported by a BBSRC Future Leader Fellowship [BB/PO11365/1] and a NIHR Southampton Biomedical Research Centre Senior Research Fellowship. We thank Carine Fixmer, Maria Lane, Benjamin Johnson, and the nurses of the Southampton Biomedical Research Unit for their help in the collection of human samples, supported by the Wessex Clinical Research Network and the National Institute of Health Research, UK. We also thank Dr. Tammie Bishop (University of Oxford) for her technical support in IHC and Prof Sir Peter Ratcliffe (University of Oxford) for the FIH antibody (mouse monoclonal 162C) and the UAS-luc/GAL4DBD-HIF1αCAD binary reporter system.

---

**Supplementary figure 1.**
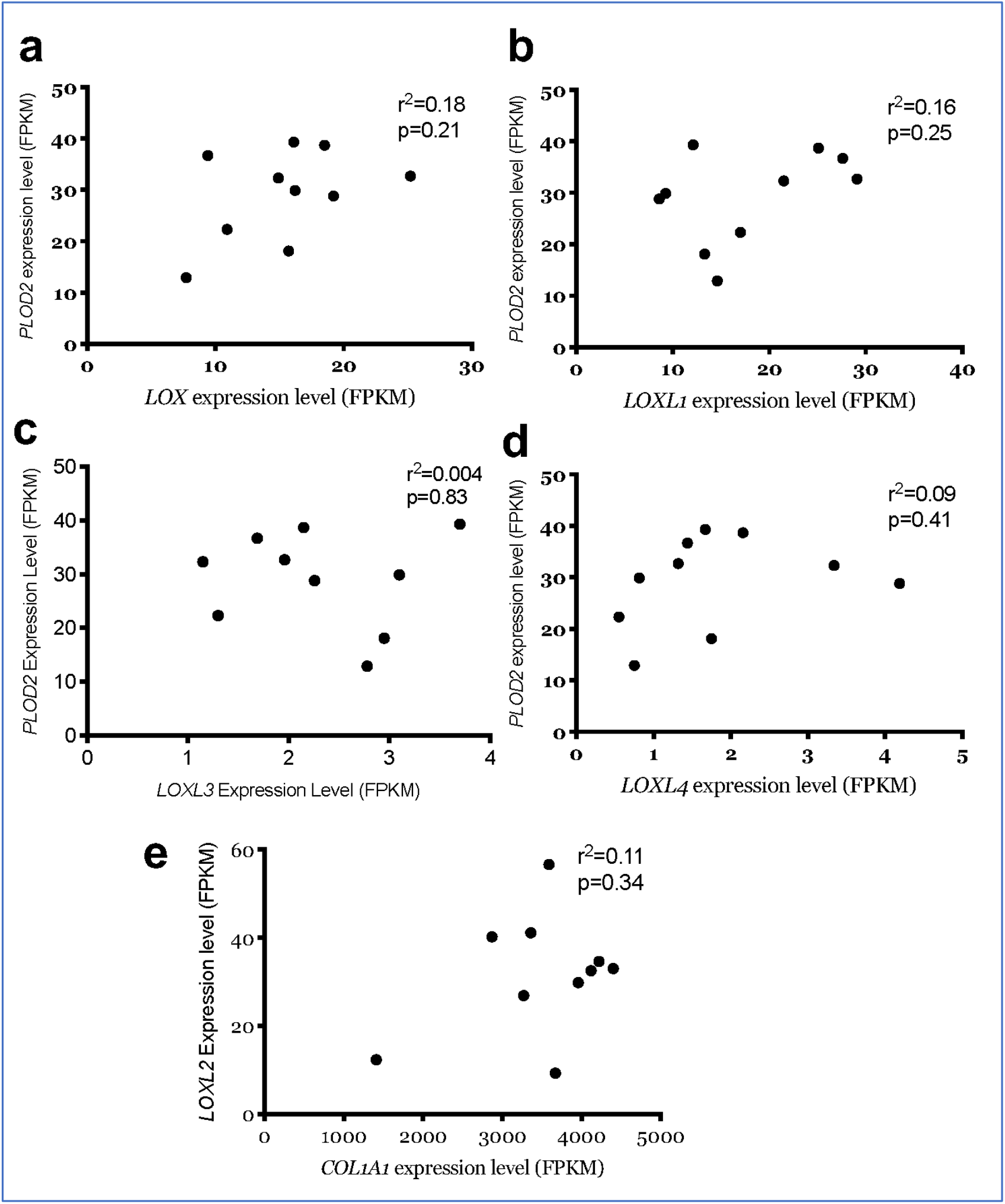
Correlation of PLOD2 with LOXL family members. (A-D) Scatterplots of paired data from (Figure 1 A-B, D-F) comparing gene expression within fibroblast foci (n=10 donors) of *PLOD2* with *LOX, LOXL1, LOXL3* and *LOXL4*. (E) Scatterplot comparing gene expression within fibroblast foci (n=10 donors) of *LOXL2* and *COL1A1* expression. Relative expression levels are calculated as Fragments Per Kilobase of transcript per Million mapped reads (FPKM). Strength of correlation calculated using Spearman correlation coefficient.

**Supplementary figure 2.**
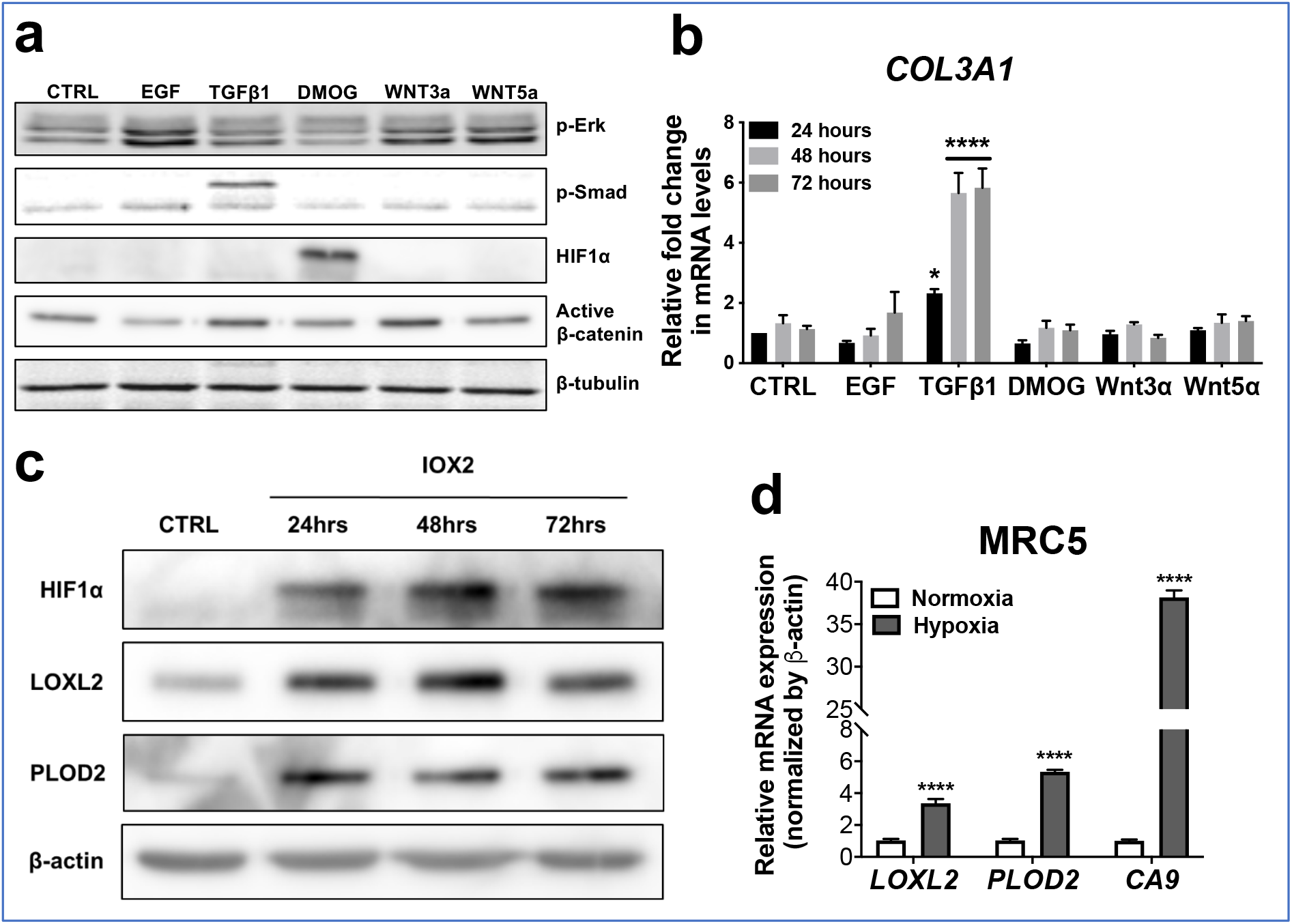
Pro-fibrotic signalling pathways in human lung fibroblasts. (A-B) Healthy lung fibroblasts exposed to control, EGF, TGFβ1, DMOG, Wnt3α or Wnt5α signalling for 24, 48 or 72 hours. n=3 independent experiments. (A) Protein expression of phosphor-ERK, phosphor-SMAD2/3, HIF1α and active β-catenin at 24 hours of exposure to conditions. β-tubulin was used as a loading control. The full blots are shown in Supplementary figure 2-source data 1. (B) Expression of COL3A1 in healthy lung fibroblasts exposed to conditions for 24, 48 or 72 hours using the ΔΔCt method. Bars indicate geometric means. ****p<0.0001 by Dunnett’s multiple comparisons test. (C) Protein expression of HIF1α, LOXL2 and PLOD2 in IPF fibroblasts exposed to control media or IOX2 for 24, 48 or 72 hours. β-actin was used as a loading control. The full blots are shown in Supplementary figure 2-source data 1. (D) Fold change in mRNA levels of *LOXL2, PLOD2* and the HIF pathway activation marker gene carbonic anhydrase IX/9 (*CA9)* in MRC5 fibroblasts after incubation in nomoxia (21% O2) or hypoxia (1% O2) for 24 hours. β-actin-normalised mRNA levels under nomoxia were used to set the baseline value at unity. Data are mean ± s.d. n = 3 samples per group. ****p<0.0001 using unpaired t test.

**Supplementary figure 3:**
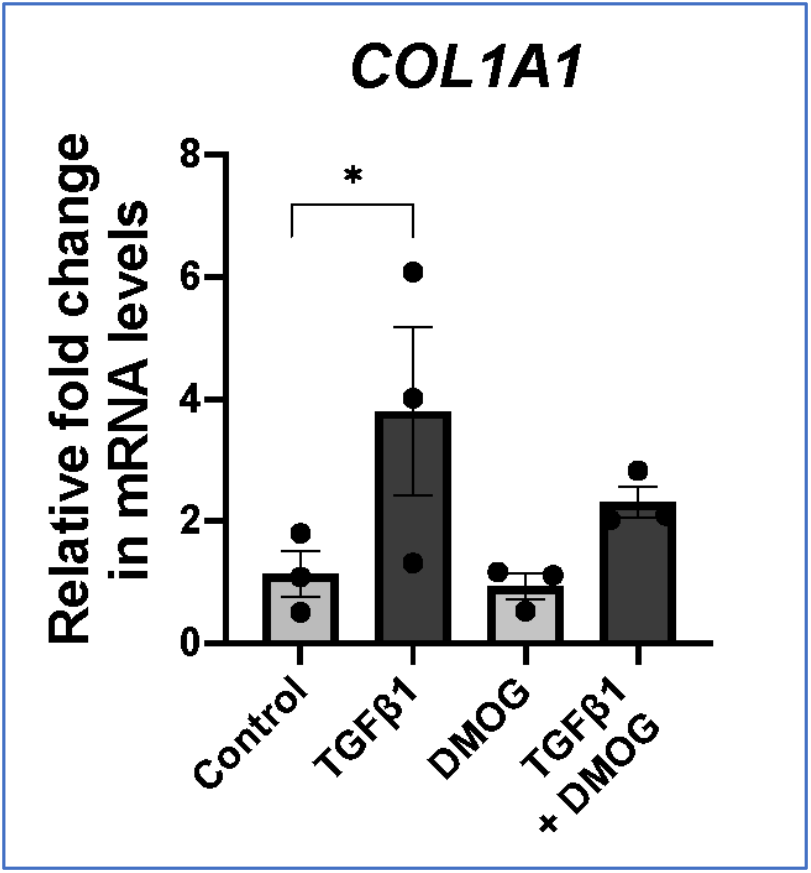
TGFβ1 promotes interstitial collagen gene expression in lung fibroblasts. Relative gene expression of *COL1A1* in lung fibroblasts from IPF donors (n=3) exposed to 48 hours of control media, TGFβ1, DMOG or combined TGFβ1 and DMOG conditions using the ΔΔCt method. Bars indicate geometric means. *p<0.05 by Dunnett’s multiple comparisons test.

**Supplementary figure 4:**
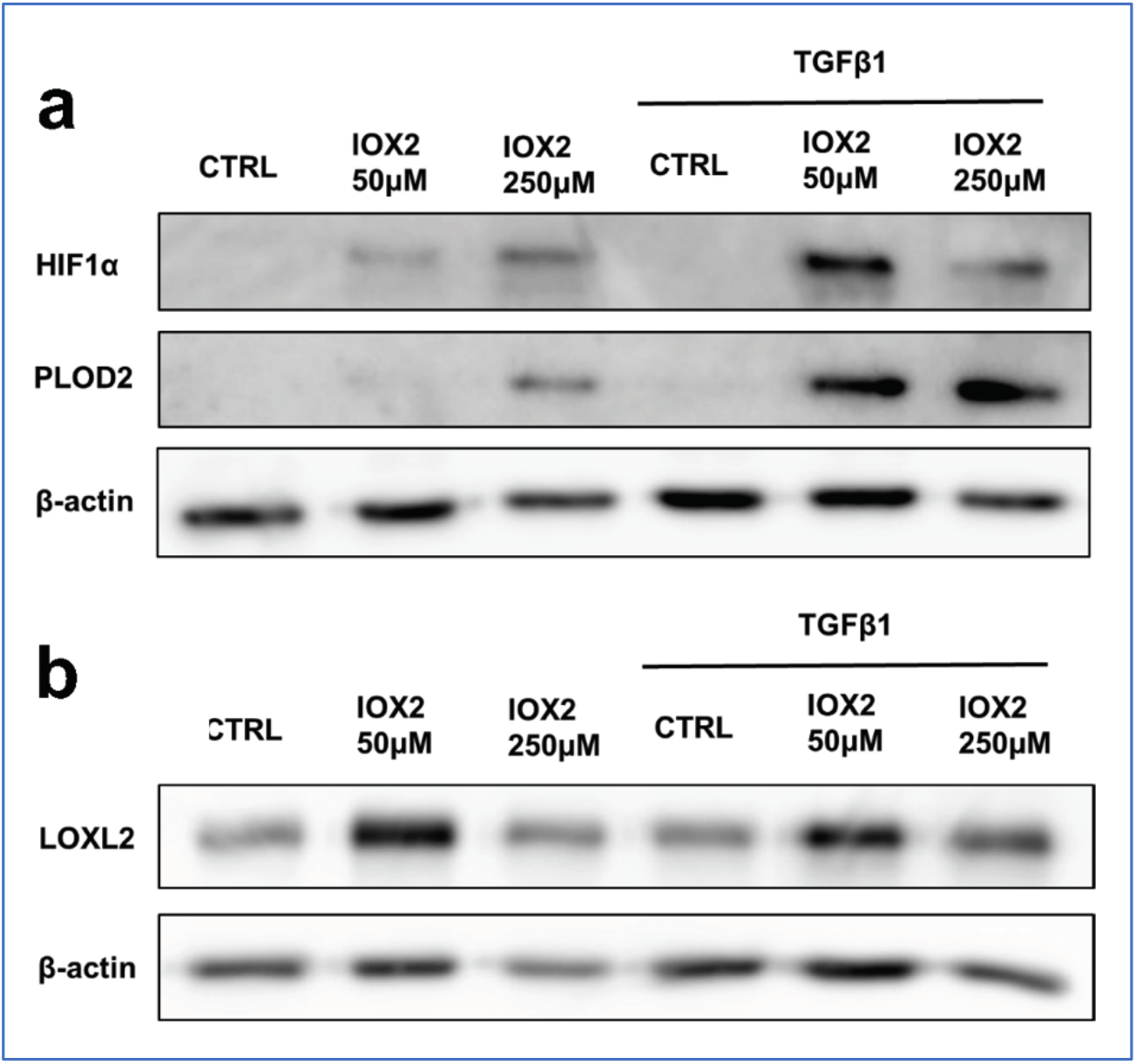
IOX2-mediated HIF pathway activation promotes PLOD2 and LOXL2 expression in the 3D *in vitro* model of fibrosis. Lung fibroblasts from IPF patients were used in the 3D model of fibrosis in the presence of IOX2 or vehicle control as indicated. Protein expression of (A) HIF1α, PLOD2 and (B) LOXL2 following two weeks of culture in the presence or absence of TGFβ1 with or without IOX2 (50 μM or 250 μM) or vehicle control. β-actin loading control. Blots representative of experiments from 2 separate IPF donors. The full blots are shown in Supplementary figure 4-source data 1.

**Supplementary figure 5:**
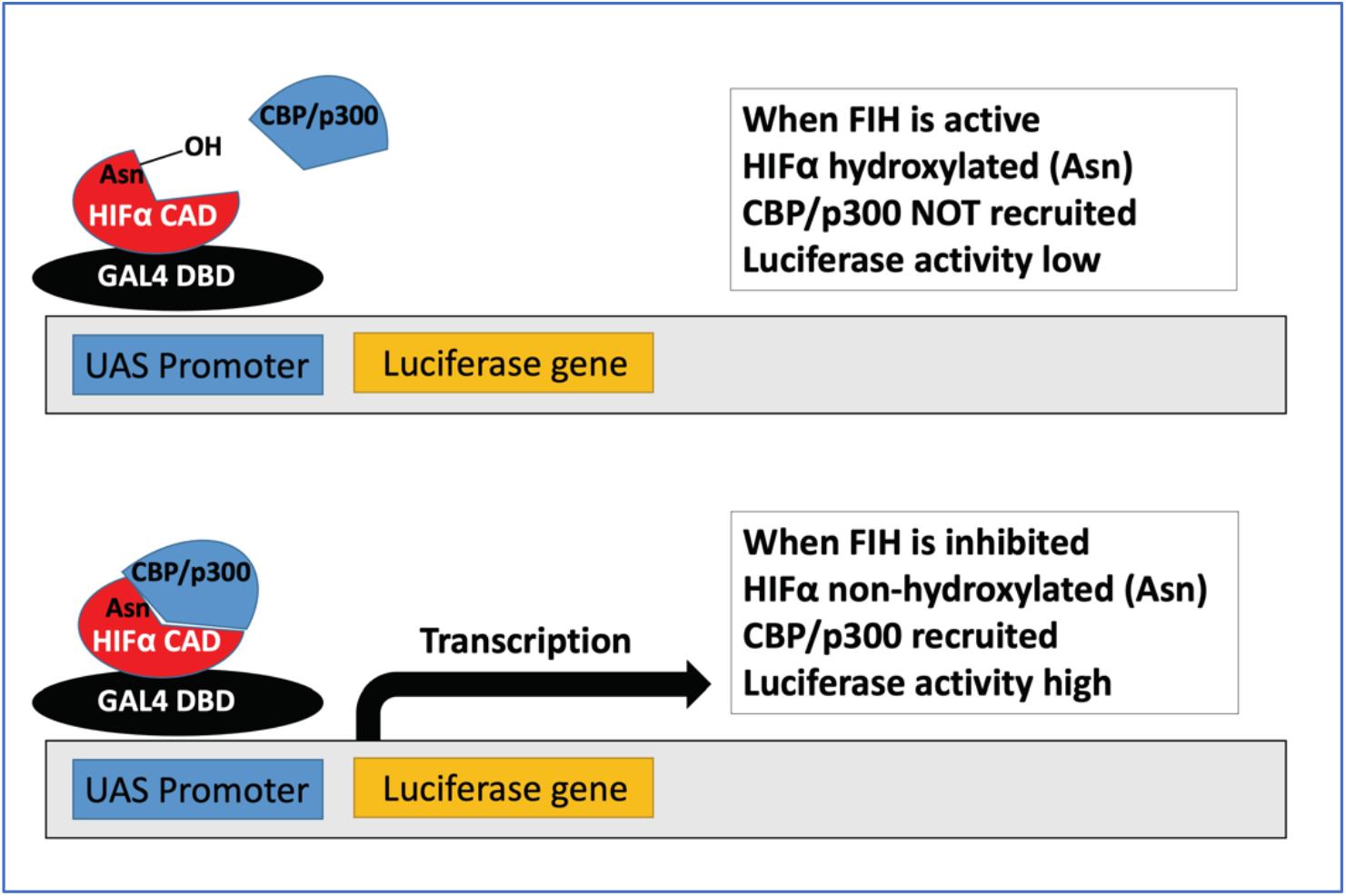
Diagram explaining the luciferase assays in Fig. 6D and E. In brief, active FIH asparaginyl hydroxylates HIF1α, inhibiting the binding of CBP/p300 and inactivating the luciferase activity. When FIH is inhibited, non-hydroxylated HIF1α binding with CBP/p300 augments luciferase activity.

**Supplementary figure 6:**
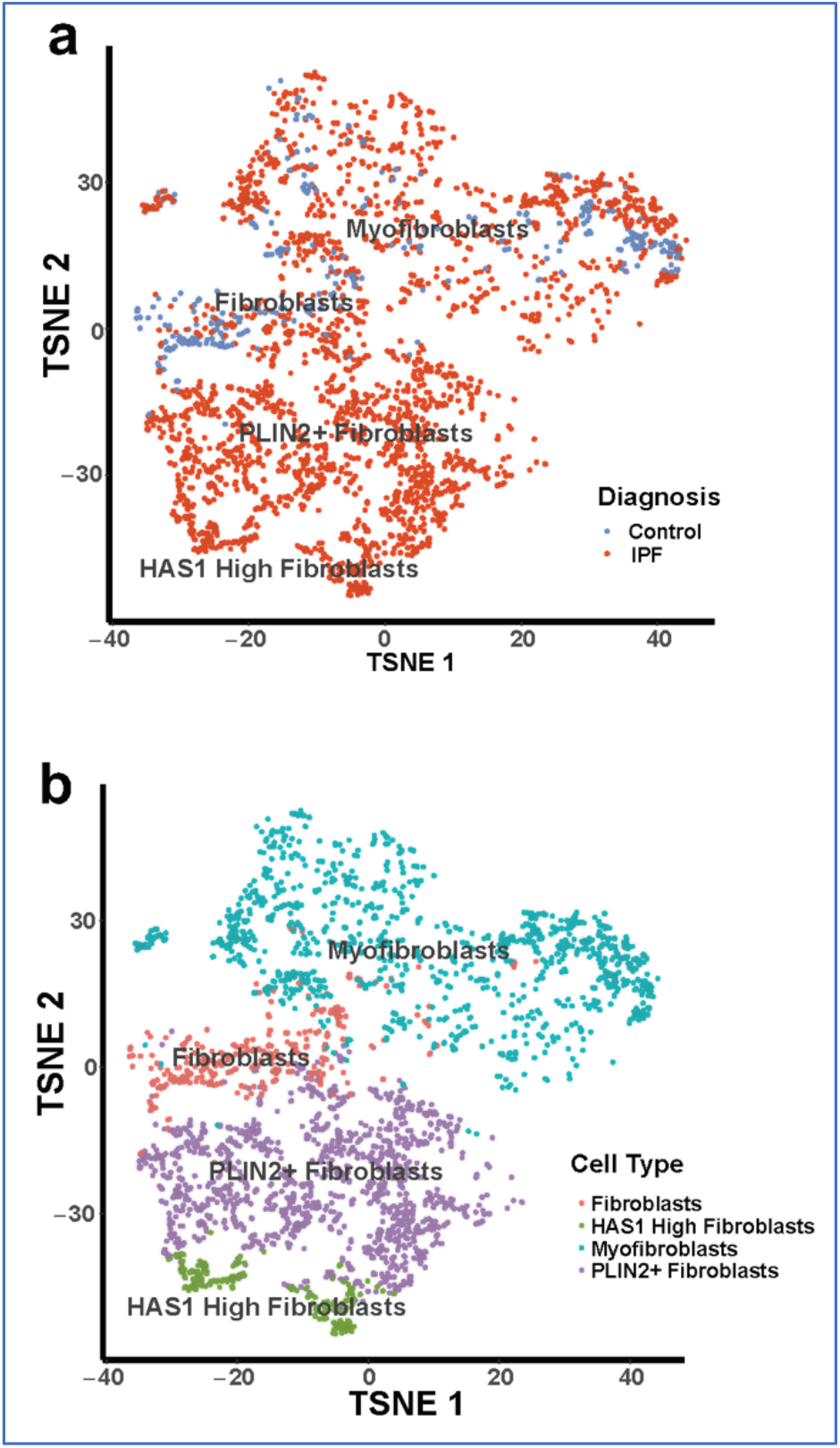
Fibroblast populations identified within a single-cell RNA sequencing dataset. (A) t-stochastic nearest neighbour embedding (t-SNE) of single cell sequencing data (GSE135893) showing clustering of different lung fibroblast types. (B) t-SNE plot of single cell fibroblast data showing diagnosis of the patients of origin for each fibroblast.

## Source Data Files

Figure 2—source data 1. Full membrane scans for western blot images for Figure 2c.

Figure 3—source data 1. Full membrane scans for western blot images for Figure 3d.

Figure 4—source data 1. Full membrane scans for western blot images for Figure 4a.

Figure 6—source data 1. Full membrane scans for western blot images for Figure 6.

Supplementary Figure 2—source data 1. Full membrane scans for western blot images for Supplementary Figure 2a,c.

Supplementary Figure 4—source data 1. Full membrane scans for western blot images for Supplementary Figure 4a,b.

Source code 1. Source code for RNAseq analyses for Figure 6a,b

Source code 2. Source code for RNAseq analyses for Figure 7a,b

Source code 3. Source code for RNAseq analyses for Figure 8

